# Serotonergic axons signal reward, sensory stimulation, and prepare for movement in primary somatosensory cortex

**DOI:** 10.64898/2026.03.19.712668

**Authors:** Patricia Przibylla, Christina Buetfering, Jakob von Engelhardt

## Abstract

Serotonin is one of the main neuromodulators in the brain, involved in regulating mood, complex behaviors and sensory input. Serotonin reaches primary somatosensory cortex (S1) via axons of neurons located in the dorsal raphe nucleus (DRN). DRN neurons can be modulated, amongst others, by reward, sensory stimulation, or movement but the activity pattern of serotonergic neurons targeting S1 is not known. Therefore, it is unclear under which circumstances serotonin is released in S1. Here, we expressed GCaMP8 in serotonergic neurons of the DRN to analyze the activity of their axons in S1 using two-photon Ca^2+^-imaging. Cluster analysis of axonal activities suggests that one to four functional groups of serotonergic axon segments project to a 0.3 mm^2^ horizontal plane of S1. We show that activity in serotonergic axons is strongly driven by reward and weakly by sensory stimulation of the whiskers. Movement, however, is preceded by a modulation, up and down, of the serotonergic signal seconds before the running onset. In summary, rewards and sensory stimulation lead to activity in serotonergic axons which is likely to adjust signal processing in S1 upon these events. The serotonergic signal changes seconds before movement onset probably preparing the neural network in S1 for the state change that accompanies running.

## Introduction

Serotonin, or 5-Hydroxytryptamin (5-HT), is a key neuromodulator in the brain. Release or presence of serotonin has been shown to influence general bodily functions such as sleep and wakefulness^1^, complex behaviors and aggression^2,3^ and also modulate sensory inputs^4-8^. When serotonin is released in primary sensory cortex (S1) it reduces the activity of principal neurons ^4,5^ and the responses to sensory stimuli^6-9^. Therefore, serotonin increases the detection threshold for sensory stimuli such as touch^7,9,10^. However, in recent years, it has been shown that S1 carries more than just sensory signals and that sensory processing is affected by events such as reward or movement^11-13^. The underlying mechanism of this modulation is not known.

Serotonergic neurons are located in the raphe nuclei of the midbrain and provide serotonin to the brain with long branching axons^14^. Serotonergic neurons projecting to S1 mostly reside in the dorsal raphe nucleus (DRN)^15^. The activity patterns of serotonergic neurons in the DRN are heterogeneous and depend on the target region of their axons^16^. Activity of DRN neurons can, for example, be regulated by wakefulness or arousal^17^, motor activity^18-20^, or reward-related behavior^18,21-24^). Whether serotonergic neurons in DRN are active in response to sensory stimulation is under debate^9,18,25^. The activity pattern of the DRN neurons branching in S1 is not known. Therefore, it is not known which behavior or event triggers serotonin release in S1 in order to modulate sensory processing.

Here, we wanted to understand under which conditions serotonergic axons are active in S1. We expressed GCaMP8^26^ in serotonergic neurons of the DRN using a SERT-Cre mouse line^27^ and imaged the activity of the axons in S1 in head-fixed behaving mice. To monitor the activity of serotonergic axons during key behaviors and events, the mouse was allowed to rest, walk or run on a running disc. In addition, we provided sensory stimulation to the whiskers and delivered sugar water rewards during the recording session. The activity of axon segments in the field of view (FOV) is functionally clustered into one to four clusters. Activity of axon segments increases after reward and sensory stimulation. Running onsets are preceded by a change in activity in serotonergic axons - up or down - with a net increase of activity just before the start of running.

## Results

### Two-photon calcium imaging of serotonergic axons in S1 in behaving mice

Serotonergic axons in S1 originate in the DRN. To record the activity of serotonergic axons in S1 we targeted the expression of GCaMP8^26^ to serotonergic neurons in the DRN. We injected an adeno-associated virus (AAV) with a flexed GCaMP8s or GCaMP8f into the DRN of SERT-Cre mice (**Fig 1A**). This limits expression of GCaMP8 to serotonergic neurons. We confirmed the labelling of serotonergic cells in the DRN and axons in S1 postmortem using immunohistochemistry in brain slices (**Fig 1B**). To record axonal activity, we implanted a chronic window above S1. Using a two-photon microscope we imaged the calcium signal of serotonergic axons in S1 (**Fig 1C, left**) in a FOV covering 577 × 575 µm^2^ (sampling rate: 40 Hz) or 556 × 556 µm^2^ (sampling rate: 20 Hz). Following registration of the axons as segments we recorded 371 +/-146 axon segments per FOV (n = 23 FOVs, 7 mice) (**Fig 1C, right**). All recordings were performed in superficial layers of S1 (75 +/-33 µm below dura, n = 23 FOVs, 7 mice) (**Fig 1D**).

**Figure 1.**
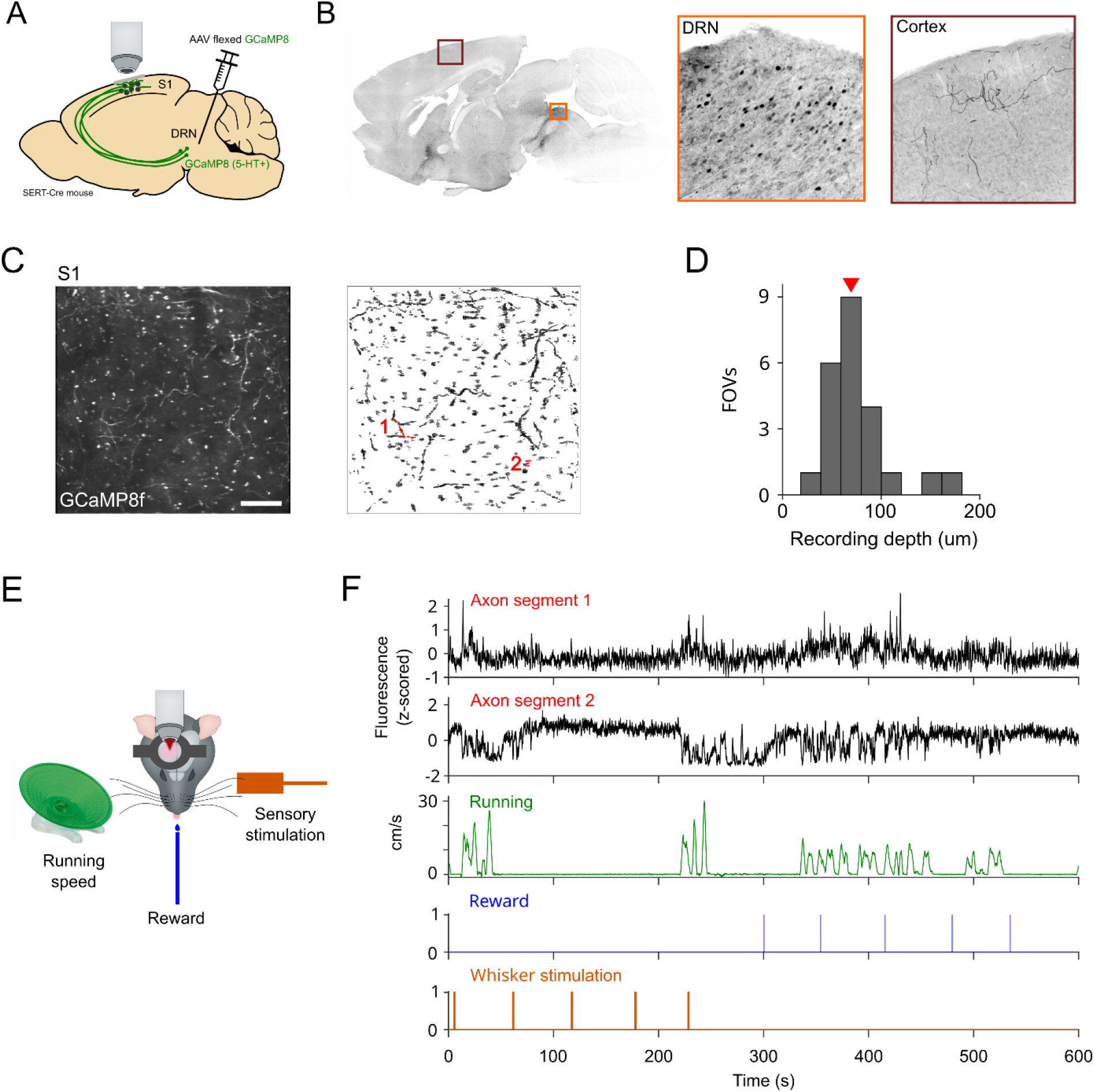
Imaging of serotonergic axons during behaviour. (A) Schematic of in vivo calcium imaging in S1 after AAV injection. Injection of AAV with flexed GCaMP8 into the dorsal raphe nucleus (DRN) of SERT-Cre mice leads to expression of GCaMP8 in serotonergic axons in primary somatosensory cortex (S1). (B) Sagittal slice with GCaMP8 staining shows the cellular labelling in the DRN and axonal labelling in S1 (left). Zoom into DRN and cortex to show serotonergic cells and axons (right). (C) Example field of view (FOV, 393 × 380 pixels) with GCaMP8f-expressing serotonergic axons (left). Scale bar: 100 µm. Registration of axon segments (right). Traces of axon segments labelled in red are shown in (F). (D) Distribution of recording depths. Red arrow indicates the recording depth of the example in (C). (E) Experimental setup under the microscope. Mice are free to run or rest on a running disc. Rewards and sensory stimulation are delivered throughout the recording session. (F) Example traces showing axonal activity of axon segments labelled in (C) as well as behavioral variables in a 10-minute recording period.

Activity of serotonergic axons was recorded in head-fixed mice that were able to run on a running disc ad libitum (**Fig 1E**). Sugar water rewards and sensory stimulation to the whisker pad with a piezo-driven pedal were given in blocks of 5 every 60 +/-10 seconds (**Fig 1E-F**). Activity of serotonergic axons was aligned to running speed (n = 23 FOVs, 7 mice), rewards and whisker stimulation (n = 16 FOVs, 5 mice) to analyse how these variables correlate with the activity of serotonergic axons (**Fig 1F**).

### Activity of axon segments in S1 is clustered

Studies in DRN have shown that serotonergic neurons have heterogeneous activity patterns^16^. In addition, they exhibit extensive branching in cortex^14^. The extent of overlap between axons of different serotonergic neurons in the cortex is unknown. Therefore, we performed pair-wise correlation analysis to quantify the similarity of activity across axon segments (**Fig 2A**). The correlation matrix was clustered using k-means clustering. Clustering was performed with increasing number of possible clusters and the clustering quality was assessed using the silhouette score^29^. For each FOV the number of clusters with the highest silhouette score was selected and included if the silhouette score was higher than 0.6 (n = 20 FOVs). Clustering results with a silhouette score lower than 0.6 are very likely not clustered (n = 3 FOVs). An example session with two clusters is shown in **Fig. 2B** after dimensionality reduction with uniform manifold approximation and projection for dimension reduction (UMAP). All sessions with valid clusters showed either 2, 3, or 4 clusters. We found that on average 2.4 +/-0.9 (n = 23 FOVs, 7 mice, mean +/-SD) functional clusters of axon segments can be found in one FOV (0.3 mm^2^) (**Fig 2C**). Each cluster can potentially contain axon segments from a single neuron in DRN or axon segments of multiple DRN neurons with similar activity preferences. We wanted to know if the axon segments of each functional group show spatial clustering, i.e. whether axon segments from the same functional group are closer to each other than to axon segments of a different functional group. We found that axon segments from a functional cluster do not cluster spatially in the FOV. The mean pairwise Euclidian distance between axon segments of the same functional cluster is similar to the pairwise distance of randomly selected axon segment pairs in the FOV (n = 52 clusters, mean pairwise Euclidian distance 223 +/-79 vs 226 +/-81 (shuffled), mean +/-SD) (**Fig 2D**). Branches from a functional group of DRN neurons therefore seem to branch widely in S1 and beyond.

**Figure 2.**
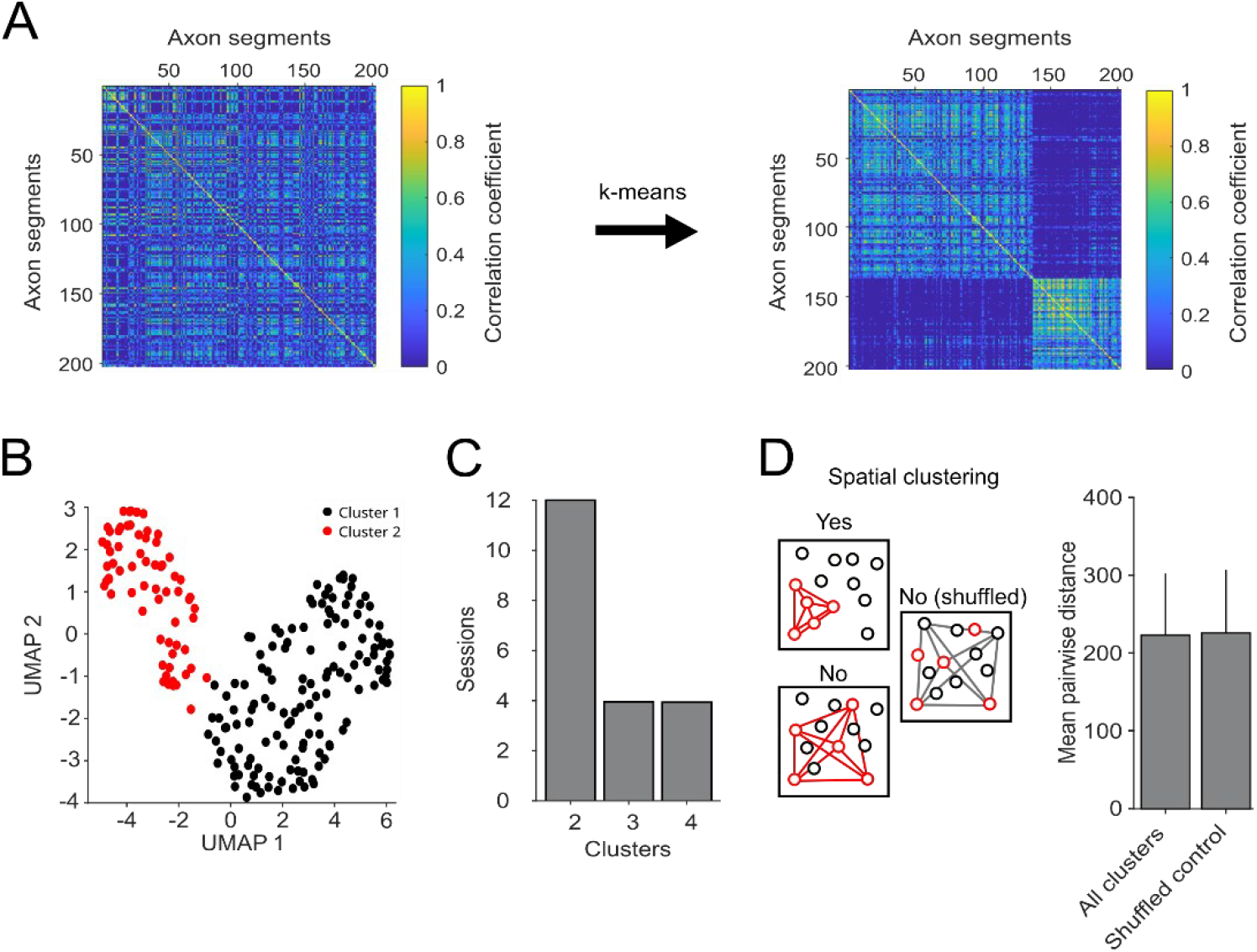
Activity of serotonergic axons is clustered. A) Pair-wise Pearson correlation of axonal activity for all axon segments in an example FOV (left). Using k-means clustering we identified functional groups of axon segments and resorted the correlation matrix (right). B) To visualise clusters, we reduced dimensionality to two dimensions using UMAP. C) For all 20 FOVs with a silhouette score above 0.6, we found that axon segments clustered into 2, 3, or 4 functional groups (n = 23 FOVs, 7 mice). The remaining 3 FOVs had a silhouette score below 0.6 and are considered not clustered. D) Schematic of analysis to test for spatial clustering (left). Mean pair-wise distances for all clusters across the 23 FOVs compared to a shuffled control (n = 52 clusters, mean +/-SD).

### Serotonergic axon segments in S1 are modulated by reward

In every recording block, five sugar water rewards were given 60 +/-10 seconds apart via a lick spout positioned in front of the snout (**Fig 3A**). Within one recording sessions (40-50 minutes), mice therefore received 20-25 rewards. **Fig 3B** shows an example axon segment with activity increasing after rewards were given. The peri-stimulus histogram (PSTH) for this axon segment shows a strong peak in activity 500 ms after the release of the reward (**Fig. 3C**). Activity then returns back to baseline within two seconds. When averaging the activity of all axon segments (n = 5171 axon segments, 16 FOVs, 5 mice, mean +/-SEM), there is a strong increase of axonal activity similar to the example axon segment (**Fig 3D**). To see whether this signal was driven by a few strongly responding segments or whether it is carried by a large majority of axon segments, we looked at the distribution of response amplitudes in standard deviation (SD) from baseline (**Fig 3E**). While most axon segments show a small increase of activity, a subset of 17 % of axon segments respond with an SD of 4 or more (n = 915 strongly-responding axon segments). Axon segments that reduced their activity after reward only make up 1% of axon segments. Splitting the axon segments based on their response amplitude, we can visualise a strong response in a subset of axons but consistent small responses in the majority of axon segments (**Fig 3F**). To exclude that this increase in activity is driven by running of mice upon receival of the reward, we extracted the running speed for the same period of time (**Fig 3G**). The running-PSTH shows now consistent onset of running upon reward receival.

**Figure 3.**
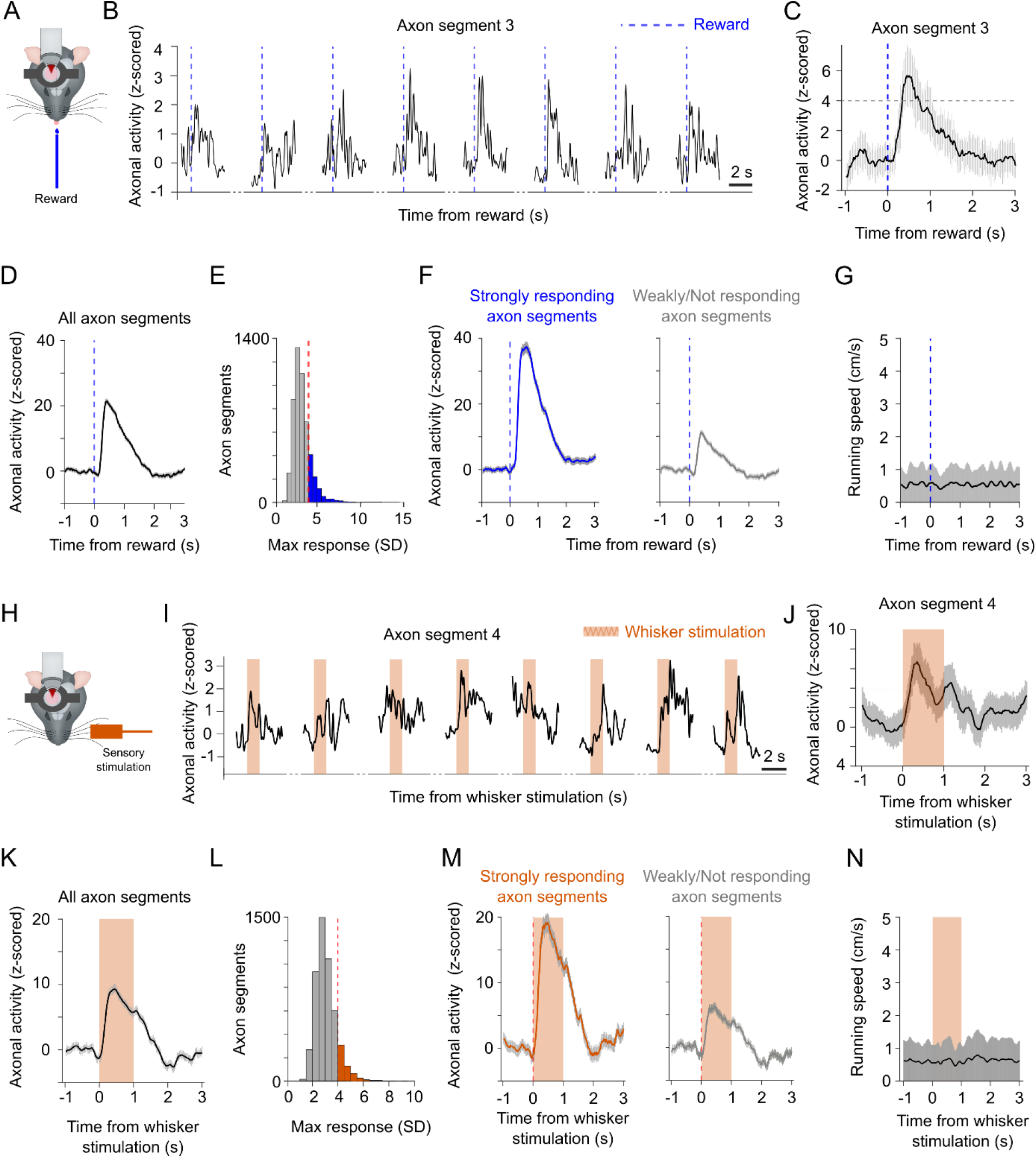
Reward and whisker stimulation trigger activity in serotonergic axons. A) Schematic of experimental setup with sugar water rewards. B) Example axon segment responding to reward. Pre- and post-reward activity shown for eight rewards in one axon segment in a recording session with a total of 20 rewards. C) Activity PSTH of example axon segment to all 20 rewards of one recording session (n = 20 rewards, mean +/-SEM). D) Activity PSTH of all axon segments to rewards (n = 5171 axon segments, 16 FOVs, 5 mice, mean +/-SEM). E) Distribution of responses to reward in standard deviation (SD) from baseline for all axon segments. All axon segments with a response larger than four SD from baseline are considered strongly responding axon segments. F) Activity PSTH of strongly (left) and weakly (right) responding axon segments aligned to reward (n = 915 strongly responding axon segments (left), n = 4229 not/weakly responding axon segments (right), 16 FOVs, 5 mice, mean +/-SEM). G) Running PSTH aligned to rewards across sessions (n = 16 FOVs, 5 mice, mean +/-SEM). H) Schematic of experimental setup with whisker stimulation. I) Example axon segment responding to whisker stimulation. Pre- and post-reward activity shown for eight whisker stimulations in a recording session with a total of 20 whisker stimulations. J) Activity PSTH of example axon segment to all 20 whisker stimulations of the recording sessions (n = 20 whisker stimulations, mean +/-SEM). K) Activity PSTH of all axon segments (n = 5171 axon segments, 16 FOVs, 5 mice, mean +/-SEM) to whisker stimulations. L) Distribution of responses to whisker stimulation in SD from baseline for all axon segments. All axon segments with a response larger than four SD from baseline are considered strongly responding axon segment. M) Activity PSTH of strongly (left) and weakly/not (right) responding axon segments aligned to reward (n = 607 strongly responding axon segments (left), n = 4527 not/weakly responding axon segments (right), 16 FOVs, 5 mice, mean +/-SEM). N) Running PSTH aligned to the whisker stimulation across sessions (n = 16 FOVs, 5 mice, mean +/-SEM).

### Serotonergic axon segments in S1 are modulated by whisker stimulation

In every recording block, five whisker stimulations were given 60 +/-10 seconds apart via a piezo-mounted pedal positioned in the whisker pad (**Fig 3H**). Within one recording sessions mice received 20-25 whisker stimulations. Each whisker stimulation lasted for 1 s with the piezo oscillating at 10 Hz. **Fig 3I** shows the activity of one axon segment that increases in activity during or after the whisker stimulation. The PSTH for this axon segment shows a strong peak in activity 300 ms after the onset of whisker stimulation. An additional small increase is seen at the offset of the whisker stimulation (**Fig 3J**). When averaging the activity of all axon segments (n = 5171 axon segments, 16 FOVs, 5 mice, mean +/-SEM), there is an increase in axonal activity with a similar temporal signature (**Fig 3K**). Axonal activity returns to baseline two seconds after the offset of the whisker stimulation. We again looked at the distribution of response amplitudes in SD to see whether the signal is carried by a majority or a minority of axon segments (**Fig 3L**). While most axon segments show a small increase of activity, a subset of 12 % of axon segments respond with an SD of 4 or more (n = 607 strongly responding axon segments) (**Fig 3M**). Axon segments that reduced their activity after whisker stimulation make up less than 1% of axon segments. To exclude that this increase in activity is driven by running of mice upon whisker stimulation, we extracted the running speed for the same period of time (n = 16 FOVs, 5 mice, mean +/-SEM) (**Fig 2N**). The PSTH of running speed shows no consistent change in running speed during whisker stimulation.

### Modulation of activity in serotonergic axon segments precedes the onset of running

Mice were allowed to run or rest on a running disc (**Fig 4A**). Example axon segments show a strong increase or decrease of their activity correlated with running (**Fig 4B**). Of note, these two axon segments were recorded from the same FOV thus some axon segments increase their activity while the same movement reduced activity in other axon segments. The activity PSTHs for these axon segments however show, that the activity does not change upon running onset but the change in axonal activity precedes the start of running (**Fig 4C**). When averaging the activity of all axon segments (n = 8532 axon segments, 23 FOVs, 7 mice, mean +/-SEM), the mean axonal activity starts to increase 1.65 s before the onset of running and decreases sharply after the onset (**Fig 4D**). All axon segments were tested for a significant increase and a significant decrease in their activity after the start of movement (**Fig 4E**). We find a large number of weakly modulated axon segments and a minority of strongly modulated axon segments. There are more strongly modulated axon segments that increase their activity before running onset (n = 1761 “UP” axon segments) than axon segments that decrease their activity (n = 876 “DOWN” axon segments). When plotting only the mean response of the strongly responding axon segments (UP and DOWN), we again see the change in axonal activity seconds before running onset (**Fig 4F**). In total, we find 18 % +/-16 % strongly up-regulated axon segments and 9 % +/-14 % strongly down-regulated axon segments (**Fig 4G**). Only a very small fraction of axon segments, i.e. 0.6 %, show a bimodal response to running onset and classify as up- and down-regulated axon segments.

**Figure 4.**
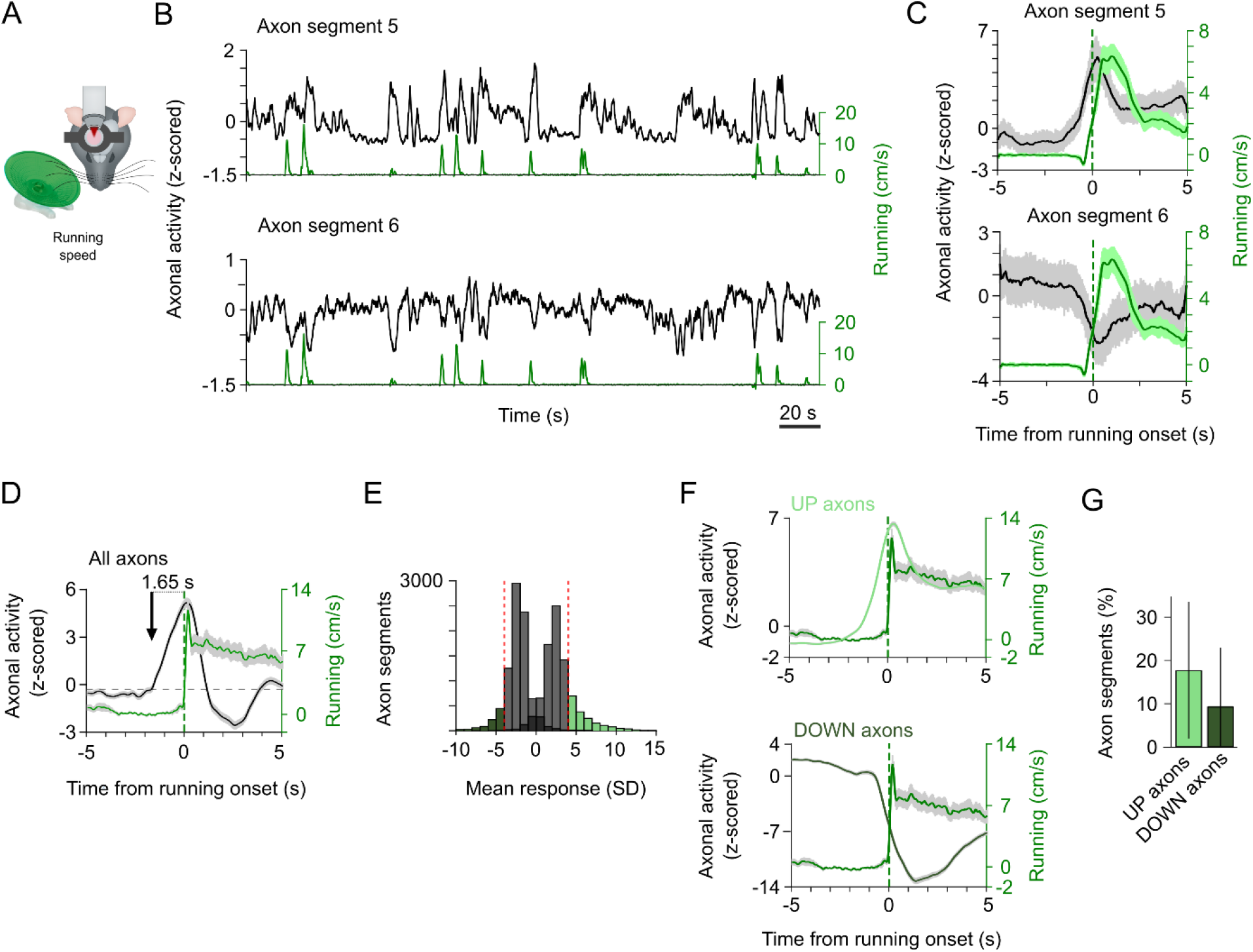
Running onset is preceded by changes in activity in serotonergic axons. A) Schematic of experimental setup to monitor running speed. B) Activity of two example axon segments from the same FOV showing positive or negative correlation to running speed. C) Activity PSTH of the two example axon segments to the running onsets in one session. D) Activity PSTH of all axon segments (n = 8532 axon segments, 23 FOVs, 7 mice, mean +/-SEM) to running onsets. Dotted line marks increase of axonal activity by 5 % from baseline. The black arrow indicates the crossing of the 5 % line indicating the rise in activity 1.65 s before running onset. E) Distribution of responses to running onsets in SD from baseline for all axon segments. All axon segments with a response larger than four SD from baseline are considered strongly responding axon segments. Strongly-responding axon segments with an increase in activity are labelled “UP axons” - axon segments with a decrease in activity are labelled “DOWN axons”. F) Top, activity PSTH of all “UP axons” aligned to running onset (n = 1761 axon segments, 23 FOVs, 7 mice, mean +/-SEM). Bottom, PSTH of all “DOWN axons” aligned to running onset (n = 876 axon segments, 23 FOVs, 7 mice, mean +/-SEM). G) Fraction of strongly responding axon segments that are increasing or decreasing their activity due to running onset across sessions (23 FOVs, 7 mice, mean +/-SD).

## Discussion

Using two-photon calcium imaging of serotonergic axons in S1 in behaving mice, we studied their activity patterns with respect to reward, sensory stimulation and running. With the aim to understand when and during what kind of events and behaviors serotonin modulates the neural circuit in S1 we recorded directly from serotonergic axons in S1. Based on known activity of DRN neurons we chose to analyse the response to two external stimuli – rewards and whisker stimulation – as well as a change in internal state, i.e. movement onset.

We found that both – reward and sensory stimulation – drive the activity in serotonergic axons with a stronger response of axons to rewards. Movement onset does not only increase activity in serotonergic axons as reported before^19,30^ but it is preceded by a change in activity of serotonergic axons of multiple seconds. This change of activity before movement is a net increase, with a sharp rise before and fall of activity directly after movement onset. The variety of signals are carried by a small number of functional groups of axons segments defined by their activity profiles through cluster analysis. Therefore, serotonin in S1 carries a reward-signal as well as a signal for sensory stimulation. Serotonin also modulates information processing in S1 before and at the onset of movement.

Whether neurons in DRN encode sensory stimuli has been the topic of debate^9,18,25^. We find a small but consistent increase of activity of serotonergic axons following whisker stimulation. We would therefore expect to find neurons in DRN that are driven by sensory input. However, in comparison to the activity modulation caused by reward or running, the response to sensory stimulation is smaller which might be the reason for the discrepancy in earlier studies. Based on data from studies showing that response amplitude in S1 reduces when serotonin is artificially released or added we would suspect that the function of serotonin release upon whisker stimulation is to avoid over-excitation of the circuit due to sensory overload or helps setting the dynamic range of sensory-evoked activity in S1^6-9^.

In comparison, reward generates a robust increase of activity in serotonergic axon activity in S1. This is in line with the finding that a subset of DRN neurons respond to reward^22,23,31^. For a long time, S1 was considered a brain area processing sensory stimuli to be broadcasted to the rest of the brain for decision making. In the last decade, many more signals have been found in S1 including decision signals, motor signals and reward-related signals^11-13^. While neurons in S1 do not seem to encode a reward signal directly, rewards shape sensory coding in S1 and reward-related information such as reward history or reward timing can be decoded from activity in S1^32,33^. The route that is used to inform S1 of reward-related activity is not clear. Feedback connections from M1 or other higher-order cortices back to S1 are a likely candidate ^34^. In humans, it has been shown that dopamine could play a role^35^. Our data show, that a reward signal in S1 is carried by serotonin released by neurons in the DRN. The neuromodulation caused by the release of serotonin would be a prime candidate to adjust sensory processing e.g. in the context of learning.

Motor activity has been linked to serotonergic signalling for over 30 years^17,19^. Seminal studies by Jacobs and colleagues show that the activity of serotonergic neurons is correlated with motor activity. They proposed that the primary function of the serotonergic system is to facilitate motor output^17^. However, the exact relationship between movement and serotonin is complex. Neurons in cat DRN, in comparison to serotonergic neurons in other raphe nuclei, do not respond to a motor challenge^36^ but some DRN neurons respond to oral-buccal movements^19^. Activity in DRN neurons can decrease during running but serotonergic axons in visual cortex increase their activity upon movement onset^20,30^. In addition, the response to serotonin appears to be context-dependent^20,37^. The impact of serotonergic neurons projecting to the spinal cord seems to be concentration-dependent and context-dependent with moderate release facilitating movement and enhancing speed and duration^20,38^. Here, we show that the activity in serotonergic axons in S1 can increase as well as decrease during locomotion and that the change of the serotonin signal starts before the onset of running. The co-existence of up- and down-regulating axon segments might generate a moderate mean release of serotonin, which – in a safe context – might translate into pro-movement modulation. Due to a faster increase and a slower decrease in the respective axon segment populations with respect to the movement onset, the net change of serotonergic modulation in S1 peaks at movement onset followed by a sharp drop. The existence of up- and down-regulating axon segments therefore could be used to sharpen the serotonin signal before and at movement onset. Serotonin would therefore act as a readiness signal or a movement-induction signal and mirror the change of the internal state. The temporally modulated change of the serotonin signal might be optimised to prepare the circuit for sensory input correlated with the running onset.

Taken together, we find that serotonin signals internal state changes and external events to S1. This is likely to allow S1 to adjust sensory processing to the context of the animal and to react to external stimuli. How exactly axonal activity modulates the neural network in S1 will be the topic of future work.

## Material and Methods

### Mice

All animal experiments were approved by the Rhineland-Palatinate State Office of Health (G22-1-031) and performed in compliance with the German Animal Welfare Act. We used SERT-Cre mice, which express Cre Recombinase under the control of the promoter of the serotonin transporter (Zhuang et al. 2005, Jax catalog no. #014554). All mice were heterozygous and had a C57Bl/6 background. Mice were bred and maintained in-house at the University Medical Center Mainz animal facility (TARC). At the start of the experiment mice were between 2-4 months old and kept in a 12-hour light-dark cycle with food and water ad libitum. Both sexes were used for the experiments.

### Surgery

Mice were anesthetized using Isoflurane (0.5–2 %) and injected with 0.1 mg kg^−1^ buprenorphine hydrochloride (Buprenovet sine, Bayer) and 10 mg kg^−1^ Carprofen (Rimadyl) 30 minutes before surgery. A small burr hole was drilled above the DRN (-4.6 AP, 0.0 ML). 500 nl of flexed GCaMP8s (CAG-FLEX-jGCaMP8s, Addgene #162380-AAV1) or GCaMP8f (CAG-FLEX-jGCaMP8f, Addgene #162382) were injected at a speed of 100 nl/min at a depth of 3 mm. A metal headplate with a 5 mm circular imaging well was fixed to the skull overlying somatosensory cortex with dental acrylic (Super-Bond C&B, Sun-Medical). A craniotomy was drilled above S1 (right hemisphere, 1.6 mm posterior and 3.5 mm lateral of bregma). A 3 mm circular glass coverslip was press-fit into the craniotomy and sealed using surgical glue (Vetbond) before fixing it with dental acrylic. Mice were allowed to recover in a warmed cage and monitored closely. At the earliest one week after the surgery habituation to the head fixation started. Mice were habituated to the lick spout and the sugar water in advance of the recordings.

### Two-photon calcium imaging

Two-photon calcium imaging was performed on a two-photon microscope (VIVO Multiphoton RS+ with Phasor, 3i). We used a ×16, 0.8 NA microscope objective (Nikon) and the Slidebook Software (3i) to record axonal activity. A Ti-Sapphire laser (Chameleon Vision, Coherent) was installed to excite GCaMP8 at 920 nm. The FOV was either 577 × 575 µm^2^ (380 × 393 pixels) with images acquired at 40 Hz or 556 × 556 µm^2^ (780 × 780 pixels) with images acquired at 20 Hz. All imaging sessions were recorded at a depth between 30 µm and 200 µm below the brain surface.

Recording behavioral variables

Axonal activity and behavioral variables were synchronized using a custom-written LABVIEW script sampling at 2kHz. To study the activity of axonal segments when rewards were given, we habituated mice to sugar water rewards (10 % sucrose solution) before the experiment. The lick spout was mounted right in front and under the snout. Rewards were delivered by opening a pinch valve controlled by a microcontroller (Arduino Uno, Arduino) for 10 ms which released 6-7 µl of sugar water. Rewards were delivered either in the first or the last 5 minutes of each recording block of 10 minutes. In a block of 5 minutes, 5 rewards were given every 60 seconds with a jitter of 20 seconds.

To study the activity of axonal segments during whisker stimulation, mice were habituated to the recording setup with a plastic pedal mounted in the middle of the left whisker pad. Each stimulation lasted 1s and the piezo oscillated at a frequency of 10 Hz in the anterior-posterior direction. The piezo (PL 140.10, PI) and its amplifier (E-650.00 LVPZT amplifier, PI) were controlled by a microcontroller (Arduino Uno, Arduino).

Running speed was measured by a rotary encoder (#140169, Kübler) mounted to the rotating center of a flat running disc (Trixie). Rotations were translated into an analog signal using a microcontroller (Arduino Mega, Arduino). All behavioral signals were fed into the LABVIEW synchronization software.

### Data analysis

ROIs corresponding to axon segments were registered and calcium signals were extracted and neuropil-corrected using Suite2P^28^. Axon segments cut horizontally as well as axons that could be traced in the imaging plane were registered as segments. Motion-correction was applied. Activity of axons was z-scored. All sessions with more than 100 axon segments were included. Data was analyzed using custom-written MATLAB scripts.

For the clustering analysis, we performed pair-wise Pearson correlation of all axon segments in a FOV. Z-scored activity was down-sampled to 1 Hz and smoothed with a gaussian filter across 10 samples. Then we used k-means clustering to partition the correlation matrix into 2-10 clusters. Cluster quality was assessed using the silhouette score. The number of clusters that reached the highest silhouette score was chosen for the FOV. All FOVs with a silhouette score larger than 0.6 were kept. A silhouette score lower than 0.6 indicates that the data does, most likely, not contain clusters^29^.

For analysis of axonal activity to reward and whisker stimulation, data was analysed at a sampling rate of 20 Hz. To assess whether axon segments respond to reward or sensory stimulation, we analysed 120 samples (3 s) after the start of the stimulus for reward and whisker stimulation. The baseline was calculated from 40 frames before the start of the stimulus. All axon segments with activity increasing more than four standard deviations from baseline were considered “strongly responding axon segments”. For plotting of peri-stimulus histograms, mean data was smoothed with a gaussian filter across 10 samples.

For the running analysis, we used data acquired at 20 or 40 Hz. All sessions recorded at 40 Hz were down-sampled to 20 Hz. We used 500 samples before running onsets to calculate the baseline. Axon segments were then tested in 200 samples (5 s) after running onset for increases and decreases in activity separately. All axon segments with samples more or less than 4 SD from baseline were considered strongly responding axon segments. For plotting, only 100 samples before and after running onset were selected and the trace was smoothed with a gaussian filter across 20 samples.

## Data availability

The data and analysis code that support the findings of this study are available from the corresponding authors upon reasonable request.

## Acknowledgment

We are grateful to B. Biesalski and A. Denner-Seckert for help with animal husbandry, R. Necel for help with equipment, and B. Grünewald for helpful discussions and comments on the manuscript. C. B. was supported by a High Potentials Grant from the University Medical Center Mainz. This work was funded by the German Research Foundation (DFG) grant INST 371/52-1, Major Research Instrumentation, 2P-Mikroscope, to J.v.E.. We thank all members and technical staff of the Institute of Pathophysiology for valuable discussions and help.

## Author contributions

C. B. and J. v. E. conceived, designed, supervised the study and acquired funding for the study. P. P. performed experiments. P. P. and C.B. analyzed the data. C. B. wrote the manuscript with critical feedback from all authors.

